# CPPVec: an accurate coding potential predictor based on a distributed representation of protein sequence

**DOI:** 10.1101/2022.05.31.494108

**Authors:** Chao Wei, Zhiwei Ye, Junying Zhang, Aimin Li

## Abstract

Long non-coding RNAs (lncRNAs) play a crucial role in numbers of biological processes and have received wide attention during the past years. Meanwhile, the rapid development of high-throughput transcriptome sequencing technologies (RNA-seq) lead to a large amount of RNA data, it is urgent to develop a fast and accurate coding potential predictor. Many computational methods have been proposed to alleviate this issue, they usually exploit information on open reading frame (ORF), k-mer, evolutionary signatures, or known protein databases. Despite the effectiveness, these methods still have much room to improve. Indeed, none of these methods exploit the context information of sequence, simple measures that are calculated with the continuous nucleotides are not enough to reflect global sequence order information. In view of this shortcoming, here, we present a novel alignment-free method, CPPVec, which exploits the global sequence order information of transcript for coding potential prediction for the first time, it can be easily implemented by distributed representation (e.g., doc2vec) of protein sequence translated from the longest ORF. Tests on human, mouse, zebrafish, fruit fly and Saccharomyces cerevisiae datasets demonstrate that CPPVec is an accurate coding potential predictor and significantly outperforms existing state-of-the-art methods.

## INTRODUCTION

Recently, long non-coding RNAs (lncRNAs, *>* 200nt) receive more and more attention for their participation in numbers of important biological processes (e.g., gene regulation and expression (1), cell cycle regulation (2)). The mutations and dysregulations in lncRNAs can cause human diseases (3), such as cancers. It is still a challenging task to discriminate lncRNAs from mRNAs, this is because 1) they often have very similar features, such as poly(A) tails, splicing and approximate sequence length (4); 2) lncRNAs may contain small open reading frame (sORF) that encodes micropeptides (5), which could induce false positives; 3) there are considerable indel errors (6) during the process of sequencing and assembly.

Many computational methods have been proposed to discriminate lncRNAs from mRNAs in the past years (7, 8, 9, 10). These methods mainly exploit five kinds of information: 1) open reading frame (ORF). The longest ORF of an RNA sequence is often extracted because it is likely to be the correct ORF where a protein is translated (11), then the ORF length, ORF integrity and ORF coverage are selected as ORF features that are effective and widely used by current methods. CPAT indicates that ORF length is the most important feature for coding potential prediction. However, ORF features are more likely to be correct when no sequencing or assembly errors occur, and hence are not suitable for platforms with indel errors, e.g., Roche (454) (12). 2) protein sequence. The physicochemical properties of the protein sequence translated from the longest ORF can also carry information for coding potential prediction. CPC2 uses isoelectric point, and CPPred adds the other two properties (e.g., gravy and instability) mentioned by CPC2. 3) *k*-mer (e.g., codon usage (3-mer), hexamer usage (6-mer)). *k*-mer features are often calculated by counting the frequency of fixed-length words (*k*-mer) that occur in an RNA sequence, or using its variant, e.g., usage frequency of adjoining nucleotide triplets (ANT) in CNCI (13). *k*-mer features are effective, and even robust (overlapping *k*-mer in PLEK (7)) for coding potential prediction for the fact that the distribution over *k*-mer is significantly different in mRNAs to lncRNAs. Despite the effectiveness, they are too short to reflect global sequence order information of RNA sequence, e.g., the position information of *k*-mer. Moreover, the increase of *k* leads to a very long and sparse feature vector, which not only induce noise, but also computational burden in real cases (14). 4) evolutionary signatures. This information is based on the sequence conservation that RNAs belongs to the same class often have similar sequence composition (e.g., base composition, transition, motifs) during the evolutionary process. CONC (15) uses amino acid composition and sequence entropy. CPPred employs CTD (composition (C), transition (T) and distribution (D)) features (16), they indicate that CTD features are particularly important for coding potential prediction of sORF. However, evolutionary signatures that these methods use are also simple statistics between the continuous nucleotides, and lose global sequence order information. 5) known protein database. As for alignment-based methods using known protein database, it is often computational expensive and also not suitable for species without annotation reference genome (7).

Based on the above analysis, we here explore how to exploit the global sequence order information of RNA sequence to enhance the performance of coding potential prediction. We developed an accurate coding potential predictor, CPPVec, which exploits the global sequence order information of RNA sequence based on distributed representation (e.g., doc2vec (17)) of protein sequence translated from the longest ORF. Tests on human, mouse, zebrafish, fruit fly and Saccharomyces cerevisiae datasets demonstrate that CPPVec significantly outperforms existing state-of-the-art methods. To our best knowledge, this is the first attempt to introduce distributed representation to coding potential prediction.

## MATERIALS AND METHODS

### Datasets

We here adopt the datasets strictly selected by CPPred to test our proposed model. Two models are built for coding potential prediction, Human-model and Integrated-model. For Human-model, human (*Homo sapiens*) samples are selected as training set and human, mouse (*Mus musculus*), zebrafish (*Danio rerio*), fruit fly (*Drosophila melanogaster*), S. cerevisiaeare selected as test sets. For Integrated-model, samples across many species, including human, mouse, zebrafish, fruit fly, S. cerevisiae, nematode (*Caenorhabditis elegans*) and thale cress (*Arabidopsis thaliana*) are selected as training set and test set. CD-hit (18) is used to remove redundancy between the test set and training set for two models. The details can be found in (10).

### Distributed representation of protein sequence

Representation learning plays an important role in machine learning methods (19). A proper representation usually achieves good result for a machine learning task. In the past years, distributed representation has been proved to be a successful data representation approach in natural language processing. Compared with one-hot encoding, distributed representation contains more semantic information about language context and more suitable for tasks such as sentiment classification (20), text classification (21). Indeed, biological sequences (e.g., DNA, RNA and protein sequences) have many similar characteristics with natural language. For one thing, they are both symbol sequences that elements in the sequence are arranged in a specified order, on the other hand, they contain a lot of semantic information, many biologists believe that biological sequences are not merely one-dimensional string of symbols, but encode a lot of useful information about molecular structure and functions in themselves (22). Hence, it is a natural idea to introduce distributed representation in natural language processing to biological sequence analysis. It is firstly introduced by (23) to protein family classification and a prediction accuracy of 99% is achieved, then it is pervasive in a wide range of applications for biological sequences analysis, e.g., protein secondary structure prediction (24), RNA-protein binding sites prediction (25, 26).

In this paper, we introduce the distributed representation to coding potential prediction for RNA sequence. To attain this goal, we are faced with three problems: 1) which kind of sequence should we choose to encode, RNA sequence, the longest ORF extracted from RNA sequence, or protein sequence extracted from the longest ORF? 2) how to build a corpus from the chosen sequences? and 3) how to train the corpus and get a distributed representation for each sequence? In our opinion, our application is concerned with coding potential of RNA sequence, and hence we should pay more attention to protein sequence extracted from RNA sequence. Moreover, just as a word in natural language, the basic unit of a protein is “word” called codon (corresponding to acid amine), and hence we consider the distributed representation of protein sequence extracted from the longest ORF, we employ the popular framework, doc2vec to generate a feature vector (embedding) of an RNA sequence. To be specific, for all the translated protein sequences, we first adopt the following splitting strategy to generate a “document” for each protein sequence:

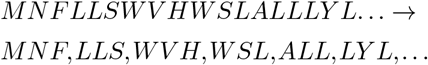

where a protein sequence is split in a non-overlapping manner with word length of 3. Second, a corpus is built from all the “documents” and trained with the distributed memory model of paragraph vectors (PV-DM), and finally the feature vector (embedding) of each protein sequence is generated and used to train a SVM classifier for coding potential prediction.

It is worth noting that in our recent paper (27), we use one-hot encoding to capture global sequence order information of biological sequence for protein coding regions prediction, however, it is not suitable for coding potential prediction for two reasons: 1) protein sequence translated from the longest ORF has a variable length but most of machine learning methods only receive a fixed-length input. 2) one-hot encoding is too low-level to reflect high-level semantic information of biological sequence. Distributed representation elegantly alleviates the above problems, e.g., doc2vec not only naturally converts a variable-length sequence to a fixed-length vector, but also contains a lot of context information of RNA sequence.

### Performance evaluation of CPPVec

To evaluate the performance of CPPVec, we use the standard performance metrics, such as sensitivity (SN), specificity (SP), accuracy (ACC), precision (PRE), F-score, AUC and MCC. These metrics can be calculated as follows:

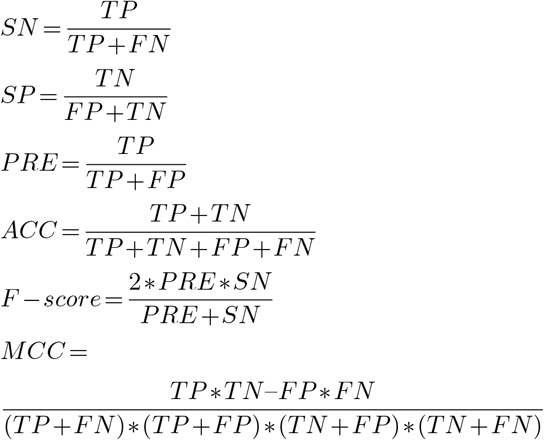

All the above metrics are based on the notions of TP, FP, TN, and FN, which correspond to number of true positives, false positives, true negatives, and false negatives. The MCC is an overall measurement of performance and another objective assessment index. AUC is the area under the receiver operating characteristic curve, it can be calculated by using the trapezoidal areas created between each ROC points.

## RESULTS AND DISCUSSION

### Pipeline of CPPVec

We used the libsvm (28) for predicting mRNAs and lncRNAs based on 138 dimensional feature vector (see Figure 1), including 100 dimensional distributed feature vector trained from protein sequence using doc2vec, and 38 features used in CPPred. All the above features are fed into a SVM classifier for coding potential prediction. The radial basis function is selected as the kernel function. The parameters of C=300 and gamma =0.4.

**Figure 1.**
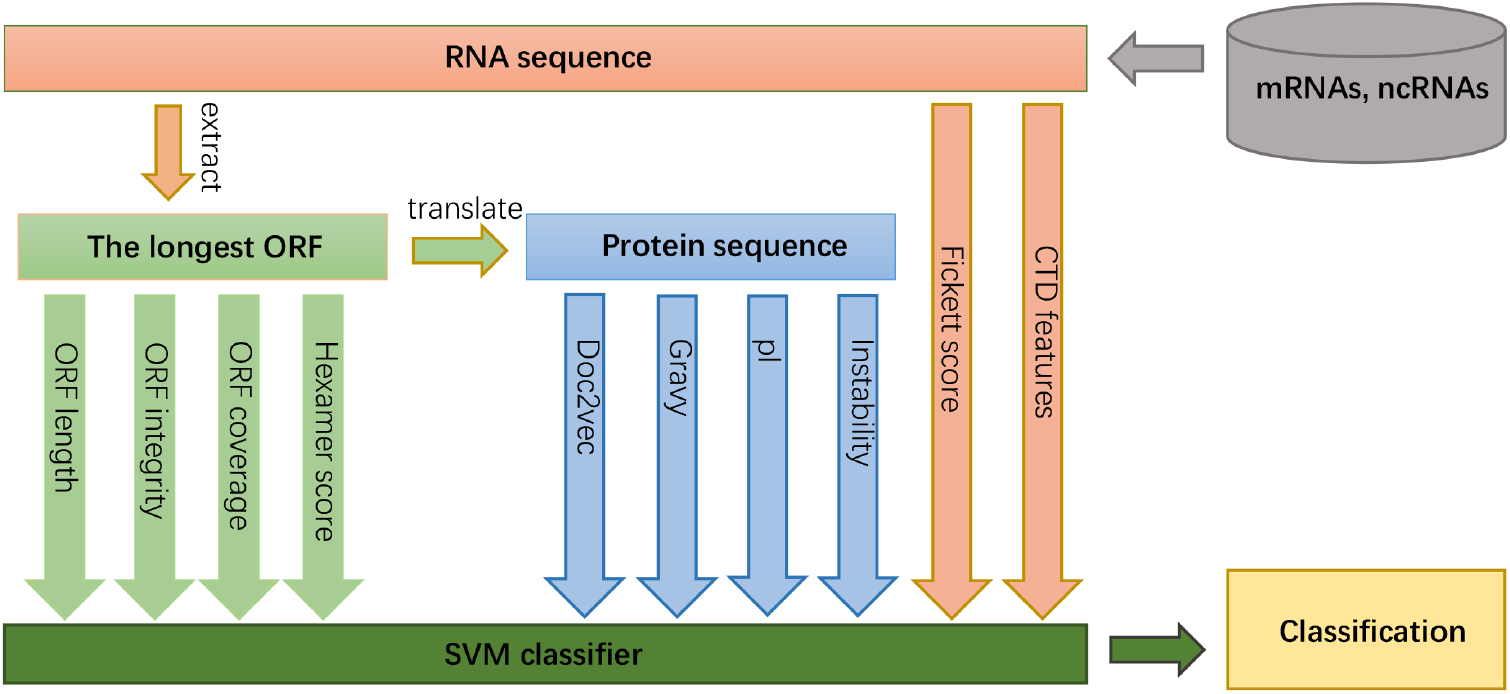
Pipeline of CPPVec. Multiple features are extracted from three kinds of sequence: the RNA sequence, the longest ORF extracted from the RNA sequence, and protein sequence translated from the longest ORF, and finally integrated into a SVM classifier for coding potential prediction. Note that the difference between CPPVec and CPPred lies in that the additional feature of doc2vec and the fixed feature of hexamer score.

### Performance of CPPVec

In order to verify the effectiveness of our proposed method, we compared our proposed method, CPPVec, with existing state-of-the-art methods, including CPPred, CPAT, CPC2, and PLEK. All the methods are trained and tested with the same datasets used in CPPred for a fair comparison. Human-model is test on human, mouse, zebrafish, S. cerevisiae and fruit fly and Integrated-model is test on Integrated-Testing.

From Table 1 to 3, it is observed that CPPVec performs the best among the existing state-of-the-art methods on all the test datasets. The MCCs of CPPVec are 0.953, 0.972 and 0.961 on Human-Testing, Mouse-Testing and Integrated-Testing, respectively, an improvement of 0.018 over the second best result achieved by PLEK on Human-Testing, 0.046 over the second best results achieved by CPPred on Mouse-Testing and 0.042 over the second best result achieved by CPPred on Integrated-Test, respectively. Moreover, CPPVec also achieved consistent results when testing with zebrafish, S. cerevisiae and fruit fly (Supplementary Tables S1–3).

**Table 1.** Comparison of CPPVec (Human-Model) and CPPred, CPAT, CPC2, PLEK on Human-Testing

| Method | SP (%) | SN (%) | PRE (%) | ACC (%) | F-score | AUC | MCC |
| --- | --- | --- | --- | --- | --- | --- | --- |
| PLEK | 98.10 | 95.42 | 98.11 | 96.73 | 0.967 | 0.993 | 0.935 |
| CPC2 | 95.30 | 90.92 | 95.26 | 93.07 | 0.930 | 0.982 | 0.862 |
| CPAT | 94.07 | 94.58 | 94.30 | 94.33 | 0.944 | 0.984 | 0.887 |
| CPPred | 97.04 | 95.44 | 97.10 | 96.23 | 0.963 | 0.992 | 0.925 |
| CPPVec | **98.69** | **96.67** | **98.71** | **97.65** | **0.977** | **0.997** | **0.953** |

**Table 2.**
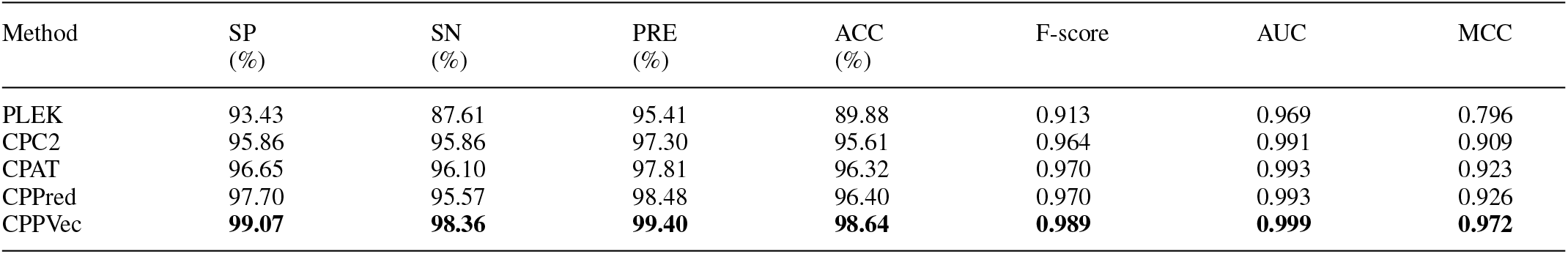
Comparison of CPPVec (Human-Model) and CPPred, CPAT, CPC2, PLEK on Mouse-Testing

| Method | SP (%) | SN (%) | PRE (%) | ACC (%) | F-score | AUC | MCC |
| --- | --- | --- | --- | --- | --- | --- | --- |
| PLEK | 93.43 | 87.61 | 95.41 | 89.88 | 0.913 | 0.969 | 0.796 |
| CPC2 | 95.86 | 95.86 | 97.30 | 95.61 | 0.964 | 0.991 | 0.909 |
| CPAT | 96.65 | 96.10 | 97.81 | 96.32 | 0.970 | 0.993 | 0.923 |
| CPPred | 97.70 | 95.57 | 98.48 | 96.40 | 0.970 | 0.993 | 0.926 |
| CPPVec | **99.07** | **98.36** | **99.40** | **98.64** | **0.989** | **0.999** | **0.972** |

**Table 3.** Comparison of CPPVec (Integrated-Model) and CPPred, CPAT, CPC2, PLEK on Integrated-Testing

| Method | SP (%) | SN (%) | PRE (%) | ACC (%) | F-score | AUC | MCC |
| --- | --- | --- | --- | --- | --- | --- | --- |
| PLEK | 90.17 | 66.32 | 87.09 | 78.24 | 0.753 | 0.872 | 0.582 |
| CPC2 | 95.54 | 91.27 | 95.34 | 93.40 | 0.933 | 0.979 | 0.869 |
| CPAT | 93.86 | 92.66 | 93.75 | 93.26 | 0.932 | 0.980 | 0.865 |
| CPPred | 94.93 | 96.91 | 95.03 | 95.92 | 0.960 | 0.990 | 0.919 |
| CPPVec | **98.38** | **97.70** | **98.38** | **98.05** | **0.981** | **0.997** | **0.961** |

### Performance of distributed representation

In order to prove the effectiveness of distributed representation of protein sequence extracted from RNA sequence, we conducted an ablation study to separate the features used in CPPVec and observe the performance improvement that distributed feature vector contributes. To be specific, we use OVEC to denote the method that only use the 100 dimensional feature vector generated from doc2vec, we use NVEC to denote the method that use features of CPPVec except distributed features. All the methods are trained on Integrated-training and test on Integrated-test. As shown in Supplementary Table S4, OVEC achieves MCC with 0.925, which even outperform CPPred that use multiple features. This result demonstrates that distributed representation of protein sequence is effective for coding potential prediction.

### Performance of fixed hexamer score

Note that in CPPred, the hexamer score is calculated with the first reading frame of the longest ORF instead of RNA sequence in CPPred. We fixed this feature for the fact that the first reading frame of the longest ORF is likely to be the correct reading frame (11).

In order to prove the effectiveness of fixed hexamer score, we compared the prediction performance of NVEC and CPPred on Integrated-Test to observe the performance improvement of fixed hexamer score. From Supplementary Table S4, NVEC shows much better prediction performance than CPPred with MCC of 0.935 versus 0.919, which proves the significance of fixed hexamer score.

## CONCLUSION

In this paper, we proposed a novel coding potential predictor (CPPVec) based on a distributed representation (e.g., doc2vec) of protein sequence translated from the longest ORF of RNA sequence, which effectively exploit the global sequence order information of protein sequence. Tests on human, mouse, fruit fly, zebrafish and S.cerevisiae demonstrates that CPPVec consistently outperforms existing state-of-the-art methods, which proves that distributed representation of protein sequence is an effective feature for coding potential prediction.

## ACKNOWLEDGEMENTS

Text. Text. Text. Text. Text. Text. Text. Text. Text. Text. Text. Text. Text. Text. Text.

## REFERENCES

1. Mercer TR, Dinger ME, Mattick JS. (2009) Long non-coding RNAs: insights into functions. Nature Reviews Genetics, 10(3), 155–159.

2. Wang X, Arai S, Song X, et al. (2008) Induced ncRNAs allosterically modify RNA-binding proteins in cis to inhibit transcription. Nature, 454(7200), 126.

3. Wapinski, O., and H. Y. Chang. (2011) Long noncoding rnas and human disease. Trends in Cell Biology, 21(6), 354–361.

4. Ulitsky, I, Bartel, et al. (2013) lincRNAs: Genomics, Evolution, and Mechanisms. Cell, 154(1), 26–46.

5. Magny E G, Pueyo J I, Pearl F, et al. (2013) Conserved regulation of cardiac calcium uptake by peptides encoded in small open reading frames. Science, 341(6150), 1116–1120.

6. Loman N J, Misra R V, Dallman T J, et al. (2013) Performance comparison of benchtop high-throughput sequencing platforms. Nature Biotechnology, 30(5), 434–439.

7. Li A, Zhang J, Zhou Z. (2014) PLEK: a tool for predicting long non-coding RNAs and messenger RNAs based on an improved k-mer scheme. BMC bioinformatics, 15(1), 1–10.

8. Kang Y J, Yang D C, Kong L, et al. (2017) CPC2: a fast and accurate coding potential calculator based on sequence intrinsic features. Nucleic acids research, 45(W1), W12–W16.

9. Wang L, Park H J, Dasari S, et al. (2013) CPAT: Coding-Potential Assessment Tool using an alignment-free logistic regression model. Nucleic acids research, 41(6), e74.

10. Tong X, Liu S. (2019) CPPred: coding potential prediction based on the global description of RNA sequence. Nucleic acids research, 47(8), e43.

11. Furuno M, Kasukawa T, Saito R, et al. (2003) CDS annotation in full-length cDNA sequence. Genome Research, 13(6B), 1478–1487.

12. Meyer M, Stenzel U, Hofreiter M. (2008) Parallel tagged sequencing on the 454 platform. Nature Protocols, 3(2), 267–278.

13. Sun L, Luo H, Bu D, et al. (2013) Utilizing sequence intrinsic composition to classify protein-coding and long non-coding transcripts. Nucleic acids research, 41(17), e166.

14. Ghandi M, Lee D, Mohammad-Noori M, et al. (2014) Enhanced regulatory sequence prediction using gapped k-mer features. PLoS computational biology, 10(7), e1003711.

15. Liu J, Gough J, Rost B. (2006) Distinguishing protein-coding from non-coding RNAs through support vector machines. PLoS genetics, 2(4), e29.

16. Dubchak I, Muchnik I, Holbrook S R, et al. (2006) Prediction of protein folding class using global description of amino acid sequence. Proceedings of the National Academy of Sciences, 92(19), 8700–8704.

17. Le Q, Mikolov T. (2014) Distributed representations of sentences and documents. International conference on machine learning, 1188–1196.

18. Li W, Godzik A. (2006) Cd-hit: a fast program for clustering and comparing large sets of protein or nucleotide sequences. Bioinformatics, 22, 1658–1659.

19. Bengio Y, Courville A, Vincent P. (2013) Representation Learning: A Review and New Perspectives. IEEE transactions on pattern analysis and machine intelligence, 35(8), 1798–1828.

20. Bollegala D, Mu T, Goulermas J Y. (2015) Cross-domain sentiment classification using sentiment sensitive embeddings. IEEE Transactions on Knowledge and Data Engineering, 28(2), 398–410.

21. Stein R A, Jaques P A, Valiati J F. (2019) An analysis of hierarchical text classification using word embeddings. Information Sciences, 471, 216–232.

22. Kuriyan J, Konforti B, Wemmer D. (2012) The molecules of life: Physical and chemical principles. WW Norton and Company.

23. Asgari E, Mofrad M R K. (2015) Continuous distributed representation of biological sequences for deep proteomics and genomics. PloS one, 10(11), e0141287.

24. Asgari E, Poerner N, McHardy A C, et al. (2019) DeepPrime2Sec: deep learning for protein secondary structure prediction from the primary sequences. bioRxiv, 705426.

25. Pan X, Shen H B. (2018) Learning distributed representations of RNA sequences and its application for predicting RNA-protein binding sites with a convolutional neural network. Neurocomputing, 305, 51–58.

26. Deng L, Liu Y, Shi Y, et al. (2020) Deep neural networks for inferring binding sites of RNA-binding proteins by using distributed representations of RNA primary sequence and secondary structure. BMC genomics, 21(13), 1–10.

27. Wei C, Zhang J, Yuan X. (2022) Enhancing the prediction of protein coding regions in biological sequence via a deep learning framework with hybrid encoding. Digital Signal Processing, 123, 103430.

28. Chang C C, Lin C J. (2011) LIBSVM: a library for support vector machines. ACM transactions on intelligent systems and technology, 2(3), 1–27.

