## Supplemental Table S1-S4 for "CPPVec: an accurate coding potential predictor based on a distributed representation of protein sequence"

**Table S1.** Comparison of CPPVec (Human-Model) and CPPred, CPAT, CPC2, PLEK on Zebrafish-Testing

| Method | SP (%) | SN (%) | PRE (%) | ACC (%) | F-score | AUC | MCC |
|---|---|---|---|---|---|---|---|
| PLEK | 88.48 | 90.48 | 91.99 | 89.67 | 0.912 | 0.962 | 0.787 |
| CPC2 | 89.95 | 96.28 | 93.34 | 93.71 | 0.948 | 0.965 | 0.869 |
| CPAT | 85.53 | 98.51 | 90.87 | 93.24 | 0.945 | 0.964 | 0.862 |
| CPPred | **93.75** | 95.55 | 95.72 | 94.82 | 0.956 | 0.979 | 0.893 |
| CPPVec | 93.57 | **98.34** | **95.72** | **96.40** | **0.970** | **0.990** | **0.926** |

**Table S2.** Comparison of CPPVec (Human-Model) and CPPred, CPAT, CPC2, PLEK on S.cerevisiae-Testing

| Method | SP (%) | SN (%) | PRE (%) | ACC (%) | F-score | AUC | MCC |
|---|---|---|---|---|---|---|---|
| PLEK | 99.03 | 46.92 | 98.73 | 49.94 | 0.638 | 0.946 | 0.216 |
| CPC2 | 100 | 88.41 | 100 | 89.08 | 0.938 | 0.983 | 0.554 |
| CPAT | 100 | 83.23 | 100 | 84.20 | 0.908 | 0.969 | 0.473 |
| CPPred | 99.76 | 86.24 | 99.98 | 87.02 | 0.926 | 0.990 | 0.515 |
| CPPVec | **100** | **93.97** | **100** | **92.23** | **0.957** | **0.994** | **0.626** |

**Table S3.** Comparison of CPPVec (Human-Model) and CPPred, CPAT, CPC2, PLEK on Fruit-fly-Testing

| Method | SP (%) | SN (%) | PRE (%) | ACC (%) | F-score | AUC | MCC |
|---|---|---|---|---|---|---|---|
| PLEK | 91.53 | 83.12 | 97.66 | 84.72 | 0.898 | 0.949 | 0.633 |
| CPC2 | 94.51 | 97.11 | 98.69 | 96.61 | 0.979 | 0.991 | 0.893 |
| CPAT | **96.85** | 97.41 | **99.24** | 97.30 | 0.983 | 0.992 | 0.916 |
| CPPred | 95.85 | 93.99 | 98.97 | 94.34 | 0.964 | 0.986 | 0.837 |
| CPPVec | 94.39 | **98.60** | 98.68 | **97.80** | **0.986** | **0.994** | **0.929** |

**Table S4.** Comparison of OVEC, NVEC, CPPred and CPPVec (Integrated-Model) on Integrated-Testing.

| Method | SP (%) | SN (%) | PRE (%) | ACC (%) | F-score | AUC | MCC |
|---|---|---|---|---|---|---|---|
| OVEC | 95.98 | 96.52 | 96.00 | 96.25 | 0.963 | 0.991 | 0.925 |
| NVEC | 96.18 | 97.32 | 96.23 | 96.75 | 0.968 | 0.992 | 0.935 |
| CPPred | 94.93 | 96.91 | 95.03 | 95.92 | 0.960 | 0.990 | 0.919 |
| CPPVec | **98.38** | **97.70** | **98.38** | **98.05** | **0.981** | **0.997** | **0.961** |
